# Dynamic cortical representations of perceptual filling-in for missing acoustic rhythm

**DOI:** 10.1101/165332

**Authors:** Francisco Cervantes Constantino, Jonathan Z. Simon

## Abstract

In the phenomenon of perceptual filling-in, missing sensory information can be reconstructed via interpolation from adjacent contextual cues by what is necessarily an endogenous, not yet well understood, neural process. In this investigation, sound stimuli were chosen to allow observation of fixed cortical oscillations driven by contextual (but missing) sensory input, thus entirely reflecting endogenous neural activity. The stimulus employed was a 5 Hz frequency-modulated tone, with brief masker probes (noise bursts) occasionally added. For half the probes, the rhythmic frequency modulation was moreover removed. Listeners reported whether the tone masked by each probe was perceived as being rhythmic or not. Time-frequency analysis of neural responses obtained by magnetoencephalography (MEG) shows that for maskers without the underlying acoustic rhythm, trials where rhythm was nonetheless perceived show higher evoked sustained rhythmic power than trials for which no rhythm was reported. The results support a model in which perceptual filling-in is aided by differential co-modulations of cortical activity at rates directly relevant to human speech communication. We propose that the presence of rhythmically-modulated neural dynamics predicts the subjective experience of a rhythmically modulated sound in real time, even when the perceptual experience is not supported by corresponding sensory data.

## Introduction

The ability to overcome the problem of missing but important sensory information, such as a conversation obscured by heavy background noise, is ethologically valuable. Even when physical information may be lost entirely, restorative phenomena such as the auditory continuity illusion, phonemic restoration, and other forms of perceptual filling-in^1–3^, allow for the percept of stable hearing in natural environments. These effects have long been hypothesized to rely on the brain’s ability to conjecture a reasonable guess as to the nature of the missing fragments^1,4^. Furthermore, as has been extensively argued, predictive coding is a task well suited for cerebral cortex^5–7^ but systematic accounts of endogenous cortical mechanisms responsible for these percepts remain unspecified.

Rhythmically-modulated sounds generate steady predictable events for which disruptions and resumptions may indicate the grouping strength of dynamic perceptual streams^8,9^. If replacement of these sounds by noise may, under some circumstances, preserve the perceived rhythm in apparent continuity, how are such streams instantiated at the neural level?Rhythmic sounds drive auditory steady-state responses (aSSR) in auditory cortex and can be recorded non-invasively via magnetoencephalography (MEG)^10–12^, with responses to rhythmic rates <10 Hz being especially prominent^13–16^. To the extent to which the neural responses track the stimulus rhythm, they can be considered sparse neural representations of the modulation rate. This experimental framework was employed to investigate the cortical effects of briefly masking and removing an ongoing low-frequency rhythmic pattern. We hypothesize that for cases where perceptual restoration of the removed rhythm occurs, the neural signature of the removal is attenuated—akin to stabilization of a cortical representation, in line with perceptual grouping under dynamic continuity. This predicts that during perceptual filling-in, the dynamical evolution of a listener’s cortical response retains oscillation in synchrony with the expected but acoustically missing rhythm.

Listeners’ perception of a continuous 5 Hz rhythmic pattern during masking was probed in a two-alternative forced choice task, where the acoustic pattern may or may not have been removed with equal probability. Simultaneously obtained MEG responses were then partitioned according to both physical and perceptual conditions, using wavelet analysis to localize oscillatory responses in time and frequency. The finding of rhythmic aSSR-like responses in cases where perceptual filling-in occurs is consistent with underlying mechanisms requiring a sustained neural representation of the restored feature^2^. Importantly, it demonstrates dynamical restoration processes occurring at scales commensurate with informal speech articulation rates^17^, as well as within MEG frequency bands that reflect cortical phase-locking to the slow temporal envelope of natural stimuli^15,18^.

## Results

### Sustained neural rhythm follows acoustic rhythm in noise

Subjects listened to four blocks (~14 min each) of a 5 Hz frequency modulated (FM) rhythmic stimulus, repeatedly masked by noise probes at pseudo-random times (see *Methods*). Half of the probes replaced the underlying rhythmic FM tone with a constant frequency tone, and half instead simply masked the underlying rhythmic stimulus, here called non-rhythmic and rhythmic probes, respectively (Fig. 1A insets). Between noise masker segments, MEG responses to steady rhythmic intervals show strong aSSR, even on a per-trial basis. Noise masker segments generate strong transient onset-like responses, after which any residual phase-locked response may disappear, on average, for rhythm-absent probes but not rhythmically-driven probes (Fig. 1A). To determine whether across subjects this change results from a decrease in aSSR power, or increased temporal jitter that would reduce averaged aSSR, inter-trial phase coherence (ITPC) and power analyses were performed on single-trial and evoked data respectively (e.g. Fig. 1B). Results of inter-trial phase coherence (ITPC) analysis reveal that, within the 0.55 – 1.22 s post probe onset interval, the ITPC difference is significant across (*N*= 35) listeners (*p* < 0.001; non-parametric permutation test). Testing for evoked rhythmic power for across listeners similarly reveals a significant difference (*p*< 0.001) within the 0.56– 1.23 s post probe onset interval. Thus the dual phase and power analyses show that both decreased aSSR power and increased intertrial jitter contribute to the decrease of the neural 5 Hz component in rhythmically absent versus driven probes.

**Figure 1.**
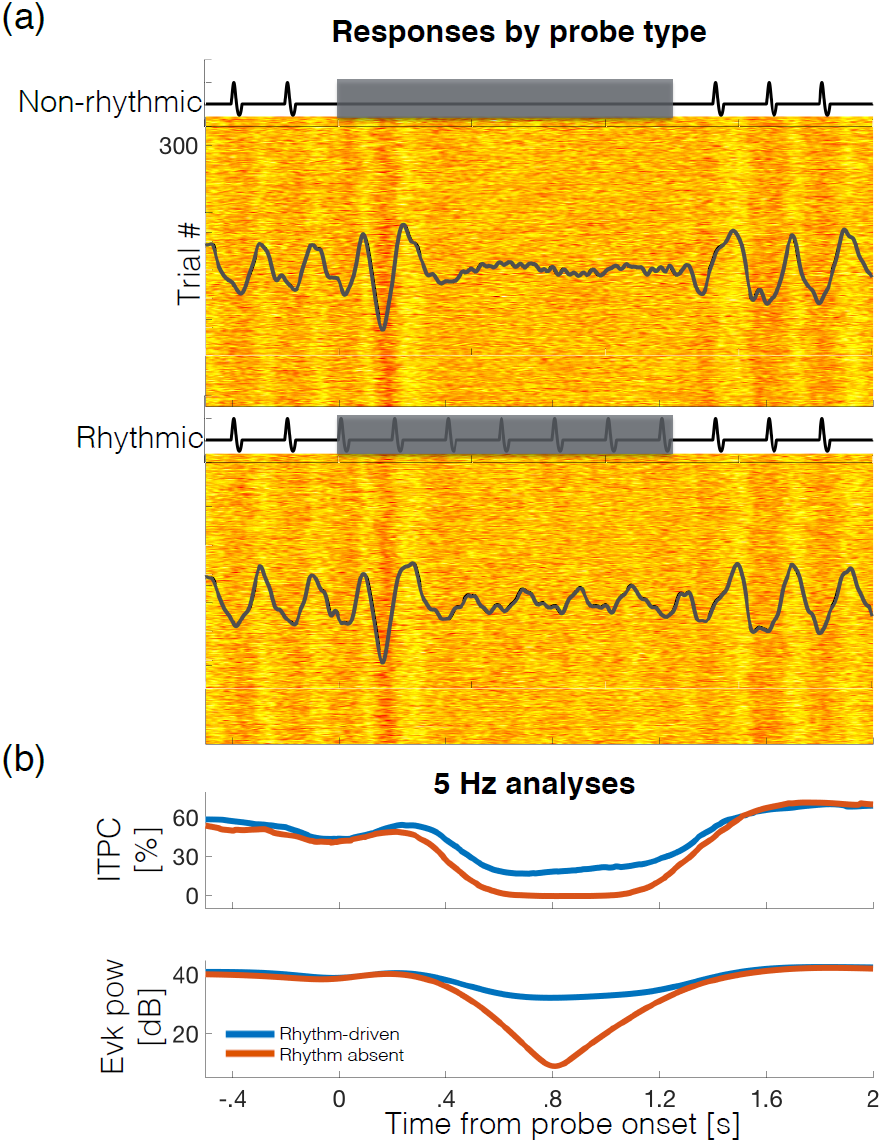
Neural representations of rhythm in noise versus tone in noise from a representative subject. (**a**) MEG responses before, during, and after a noise probe are shown (single MEG component obtained via spatial filtering; see *Methods* and Supplementary Fig. 1). The basic stimulus consists of a 5 Hz pulsatile (short duty-cycle) FM tone, centered at *f*_*0*_ = 1024 Hz, to which 1.24 s noise probes were applied. Insets: illustration of a non-rhythmic probe where pulses are replaced by the constant tone (top); and a rhythmic probe, where the FM continues under the noise (bottom). Before and after the probes, phase locking to the main rhythmic stimulus is apparent even on a per-trial basis. Overlaid on each response raster, evoked activity (averaged separately for each probe type) reveals a measurable aSSR during rhythmically-driven probes (top) but not during rhythm-absent probes (bottom). (**b**) Top: Phase analysis at 5 Hz shows estimated phase-locking over time as measured by ITPC. During masking ITPC values drop to near floor in rhythm absent probes (orange) but only to half of baseline levels in rhythm-driven probes (blue). Bottom: Analysis of spectral power (also at the 5 Hz rhythm rate) also shows considerable difference between probe types for this subject.

### Sustained neural rhythm follows listeners’ perceived rhythm in noise

In order to determine how neural representations of rhythm co-varied with perception, after each trial the probe was classified by the subject as perceived as rhythmic or as non-rhythmic. This resulted in a 2-by-2 partition of analyzed trials: (1) non-rhythmic probes perceived rhythmic (‘filling-in’); (2) non-rhythmic probes perceived non-rhythmic (rhythm ‘absent’); (3) rhythmic probes perceived rhythmic (rhythm ‘present’); and (4) rhythmic probes perceived non-rhythmic (rhythm ‘missed’). Fig. 2 shows the grand average evoked 5 Hz response power before, during, and after noise probes, for each combined condition of stimulus and percept.Transient (and broadband) masker-onset responses were evident during the initial 0.3 s post masker onset (cf. Supplementary Fig. 2) (brief pre-causal dips accompanying these transients are due to convolution residuals from the continuous wavelet transform).

**Figure 2.**
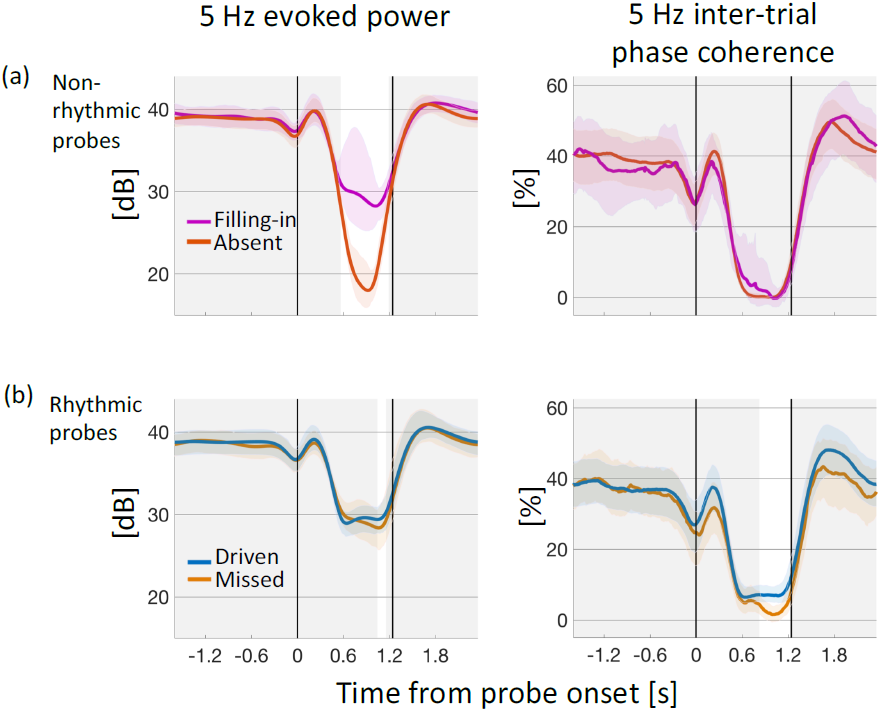
Percept-specific endogenous representations of patterned sound. Grand averages (*N* = 35) of rhythmic evoked power and intertrial phase coherence partitioned by probe type and reported percept. Noise probe starts at the first vertical line at *t* = 0 s and continues until the next vertical line at *t* = 1.24 s. (**a**) Non-rhythmic probes: (Left) After an initial transient, rhythmic evoked power was reduced regardless of percept, but differentially by 7.9 dB depending on percept as present (magenta), or absent (orange). (Right) No significant difference was observed for ITPC, where there was a reduction to near floor during the probe. (**b**) Rhythmic probes: (Left) During masking, rhythmic evoked power drops by 9.5 dB in average, holding relatively steady for the duration of the probe. (Right) Similarly, inter-trial phase coherence drops by about 81% for the duration of the probe. For probes in which the rhythm was missed (brown), however, both evoked power and ITPC showed an additional reduction (only near the end of the probe) compared to rhythmically-driven probes (blue). Solid lines: mean across subjects and trials; Color bands: bootstrap 95% confidence of the mean over subjects; Grey bands: time intervals with no significant difference by percept.

For non-rhythmic probes (Fig. 2A), phase coherence dropped to almost 0% for both perceptual conditions (filling-in and absent, right panel). Rhythmic spectral power also dropped from the initial baseline for both perceptual states, but the decrease was on average 7.9 dB worse when subjects reported the rhythm absent than present (filling-in). Decreases were restored to baseline values by 0.8 to 1.2 s post probe offset (equivalent to between 4 and 6 rhythmic pulse cycles. Thus, within non-rhythmic probes, a sustained and significant percept-specific difference was observed in rhythmic evoked power (0.56 to 1.19 s, *p* < 0.001), but this was not the case for phase locking (*p*> 0.18).

For rhythmic probes (Fig. 2B), the masker was associated with an average relative decrease of 9.5 dB evoked power regardless of perceptual condition (driven and missed), and with a relative decrease of ~75% in trial-to-trial phase locking. When subjects missed the rhythm, evoked power and inter-trial phase coherence both further decreased, with percept-specific decreases sustained over a longer period for ITPC (0.84 – 1.25 s, *p* < 0.001; right panel) than evoked power (1.04 – 1.15 s, *p* = 0.008; left panel).

### Rhythmic neural power as discrimination statistic in a rhythm detection task

With the observation that differential neural processing of masked rhythm depends on listeners’ percept, it was next investigated whether the observed divergence might have properties of an internal variable underlying discrimination. Based on the previous result, we hypothesized that the 5 Hz target neural processing power in the final ~600 ms of the probe interval might act as such variable. For each subject, a metric was created from the rhythmic evoked power differences contrast, integrated over the 0.56 – 1.24 s interval of interest post probe onset. To illustrate the use of this latent variable as a discrimination statistic, a bootstrap resampling of trials (with replacement) was used to produce distributions of evoked power sustained over the critical window (two representative subjects shown in Fig. 3A). A neural discriminability metric was then computed from their relative separation (see ***Methods*).** To assess the potential of this sustained evoked power to operate as a variable relevant to perceptual discrimination, the neural metric was compared with psychometric *d*’ scores that index behavioral sensitivity of listeners to the detection task^19^ (Fig. 3B, blue), with the result that the two are significantly correlated (*ρ* = 0.728, *p*=1.04 ×10^-6^).

**Figure 3.**
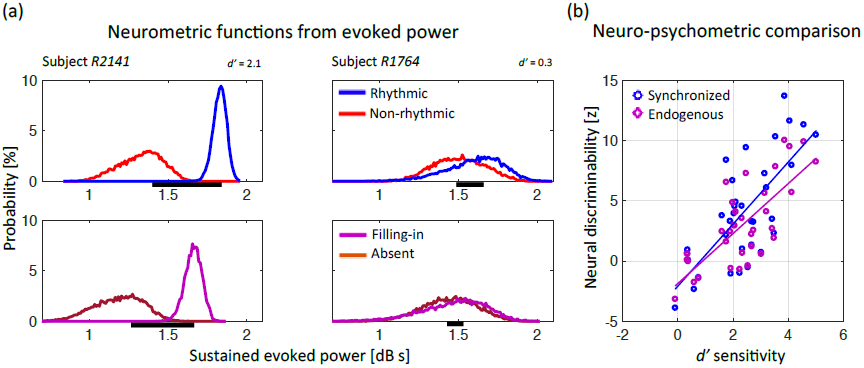
Rhythmic target power acts as a discriminant neural statistic for perceived rhythm. (**a**) Top: In two representative subjects, behavior covaries with empirically-derived neural discriminability distributions. Probability distributions of a given level of sustained (time-integrated) evoked power depend on the acoustic presence (blue) or absence (red) of stimulus rhythmic FM; a neural discriminability score (proportional to horizontal black bar length) can be obtained from them. In the first subject (left panel), the small overlap between the distributions gives high neural discriminability; for the second subject (right panel), both distributions overlap substantially, giving poor discriminability. Bottom: Next, empirically-derived neural distributions were obtained *only from non-rhythmic probes* (i.e., the red curves in the top panels), now conditioned instead by percept. A similar pattern in the distributions is observed. Distributions obtained via bootstrap. (**b**) Over subjects, the psychometric *d*’ sensitivity index (abscissa) correlates with the neurometric discriminability index based on acoustic contrast (rhythmic versus non-rhythmic probe, blue;*ρ* = 0.73, *p* = 1.0×10^-6^). Critically, behavioral sensitivity to ‘filling-in’ also correlates with rhythmic evoked power differences despite the absence of stimulus rhythm via the related neurometric discriminability index based on perceptual contrast (filling-in versus reported absent, magenta; *ρ* = 0.69, *p* = 6.1×10^-6^).

A related latent discrimination statistic, directly relevant to the phenomenon of filling-in, is computed with contributions only from endogenous (non-sensory) factors, by analyzing the responses to non-rhythmic probes exclusively(Fig. 3A, bottom). In these purely percept-specific (constant acoustics) distributions, neural power discriminability was defined analogously as the difference in rhythmic evoked power between filling-in and rhythm-absent trials, integrated over the time at which significant differences were observed at the group level in the previous section (0.56 to 1.19 s post probe onset, as in Fig. 2B). Just as for the acoustic contrasts, this discriminability index also correlates strongly with the psychometric sensitivity indices across listeners (Fig. 3B, magenta) (*ρ*=0.745, *p*=4.23×10^-7^). Thus, consistent with the properties of a latent discrimination statistic, sustained evoked power may account for both stimulus-and percept-specific differential processing, where the latter reflects only endogenous neural processes.

### Spectrum of power increase in target-related neural rhythm dynamics with filling-in

Given the possibility that increased power at the 5 Hz rhythmic frequency would be accompanied by increased spectral power at other frequencies, it is important to consider whether change arises as a power gain specific to the target frequency or as a modulatory effect over a larger spectral region that includes the target frequency band. By extending the wavelet analysis over a broader frequency range (1-25 Hz), the spectral extent of restoration was probed to address whether changes are target-specific, or instead accompanied by other activity that may be behaviorally relevant.

Evoked power analyses across probe conditions and subjects reveal that the evoked response contains two frequency ranges, one centered on the target 5 Hz, and the other centered on the 10 Hz first harmonic (Fig. 4A). To analyze time-frequency power contrast between conditions, corresponding spectrograms (baseline corrected per frequency band) were subtracted. In particular, the ‘driven’ minus ‘absent’ map results in a contrast whose differences arise from synchronization to physical differences in the sound, while ‘filling-in’minus‘absent’ maps differences due entirely to endogenous activity (Fig. 4B, left panels). For the first case, the defined ‘synchronized’ contrast (Fig. 4B, top left) group average data shows a spectrotemporal region, ~600 ms post probe offset until the end of the probe, of significant differential neural processing (*p* = 3.3×10^-4^), rooted in physical stimuli differences. The region is limited to the spectral neighborhood of the target (half maximum 4.1-6.7 Hz; maximum 3.8-7.5 Hz), which may be expected as smearing from Fourier/Heisenberg uncertainty. For the ‘endogenous’ contrast (Fig. 4B, bottom left), a similar profile was found (half maximum 4.1-6.6 Hz; maximum 3.8-6.8 Hz; *p* = 6.7×10^-4^), with additional enhancement around the target first harmonic (0.4 to 1.1 s post probe onset; half maximum 9.7-11 Hz; maximum 8.9 to 11.9 Hz; *p* = 0.01). In a related analysis of a third partition contrast, ‘rhythm-driven’ minus ‘missed’, no spectrotemporal cluster of significance was found (*p*=0.29).

**Figure 4.**
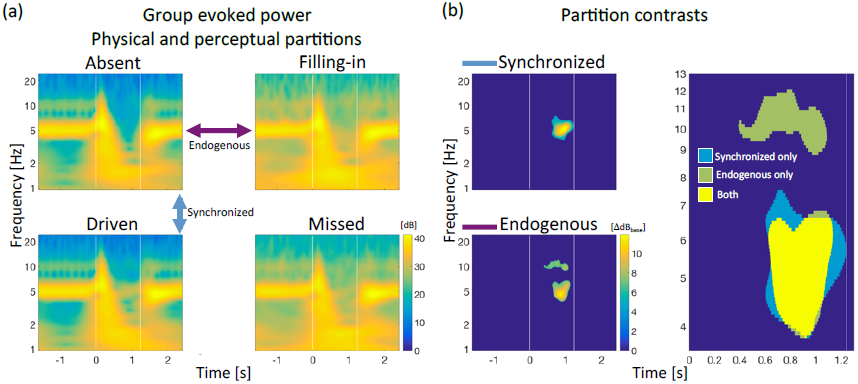
Stimulus-and percept-specific spectrotemporal modulations of cortical activity during restored rhythm. (**a**) Wavelet power correlograms, in a 1-25 Hz frequency range, reveal qualitative differences in steady neural responses post probe onset, across participants (*N* = 32). Color arrows indicate spectrogram pairs submitted to difference contrasts as follows. (**b**) Differences between spectrograms reveal differential processing under alternative percepts, whether based on different physical sounds (top left), or on endogenous restorative processes (bottom left), in both cases specific to the target 5 Hz frequency band. The latter case of filling-in generates enhanced sustained power in the first harmonic band (~10 Hz) as well. Synchronization maps are shown masked by regions of group-level significance, as determined by permutations within contrast pairs, performed independently across subjects (‘driven’, *p* = 3.3×10^-4^;‘filling-in’-near 5 Hz *p* = 6.7×10^-4^, filling-in’near10 Hz *p* = 0.01). The lower-rate rhythmic enhancements (~5 Hz) coincide spectrotemporally even though the sensory bases for each are different (right). White vertical lines indicate noise probe temporal edges.

Upon examination of whether the additional spectral information conveyed by these maps improved neural predictions regarding listeners’ behavior, we found that neural discriminability indices based upon the ‘synchronized’ region in this section showed no improvement over the target frequency specific index obtained previously for 5 Hz only measures (*ρ* = 0.53; *p* = 0.001). The ‘endogenous’ regions, jointly, showed no improvement in predictive power of listener’s performance (*ρ* = 0.72; *p* = 1.4×10^-6^) over that of the target-based index alone. Separating these regions into 5 Hz and 10 Hz domains revealed that the lower (target rhythm) region was more predictive (5 Hz only: *ρ* = 0.73, *p* = 8.2 × 10^-7^; 10 Hz only: *ρ* = 0.44, *p* = 0.01). These results suggest that differential narrowband 5 Hz power is most critical to explain listeners’ detection performance shown previously, and that for filling-in trials, some improvement also arises from integrating over the broadened filter to include neighbor target frequencies present in the average timeseries of endogenous neural activity.

## Discussion

The subjective experience of attending effectively to complex sound scenes in noisy environments can be substantially assisted by perceptual restoration. This effect is investigated using MEG to record the neural dynamics of a steady temporal pattern while repaired perceptually. Measures of differential cortical processing contributed to the identification of a discrimination statistic predicting a subject’s behavioral performance sensitivity. The data are consistent with the view that perceptual restoration is attributable to endogenous neural processes, emerging from learnable temporal patterns present in the tracked auditory object, at modulation rates that dominate natural communication speech sounds.

Perceptual restoration, the effect of hearing the continuation of a sound regardless of an interrupting masker, includes descriptions of “auditory induction”, “temporal induction”, “perceptual synthesis”, or “contextual catenation” of dynamic sounds in classic studies^20,21^. It implies an ability to discount disruptive but extraneous interruptions to relevant acoustic signals, so much so that even noise-filled *gaps* are more likely to be discounted as such^21^.Where multiple interpretations of a relevant acoustic signal are possible (e.g. phonemes), perceptual restoration has been probed in identification tasks; for more constrained decision spaces, it may be probed based on sound delivery quality assessments, such as gap localization of the excised token signal (e.g. Warren’s paradigm^21^), and by discrimination of noise-added vs. noise-replaced token gap alternatives (e.g. Samuel’s paradigm^22^). Our method subscribes to the latter approach, also referred to as ‘filling-in’, which emphasizes the signal detection strategy followed in cases where a listener classification is inconsistent with the token absence in a gap^3,23–29^. As has been noted^30^, from the listener’s utilitarian perspective, this effect of induction in a challenging environment is not aimed at the production of decision errors (or illusions) but to assist against masking. Restoration refers to the perception of a token projected by a context (such as a speaker’s intention), with apparent intactness^30^. Critical to this is a strong masker, along with contextual evidence favoring a specific acoustic token with high probability. This combination allows inference that the lack of auditory evidence of the token could be ascribed to energetic masking^1,4,31^.

A simple and compelling example of perceptual restoration is that of a pure tone followed by a brief noise-filled gap where the tone has been excised: this leads to a strong illusory percept of continuity of the tone^32^. The percept appears to rely on two related effects, the more obvious being conveying the original signal as uninterrupted, but also, critically, accompanied by an attenuation of discontinuity boundaries^33^. Neural correlates of both effects have been observed in single units in macaque primary auditory cortex (A1), where up to 35% of sampled single units respond to a gap with noise as though the tone were continuously present^34,35^. In some cases there is also failure of a transient response at the end of the gap^34^.For human listeners, there is evidence that such compensatory principles may extend to disruptions to dynamically modulated sound, including amplitude-modulated (AM) sound, single vowels, and consonants within words^8,9,24,28,36,37^, the latter of which fall under the concept of phonemic restoration^1,3,22^. Depending on stimulus, neural correlates have been localized to different areas, including Heschl’s gyrus for missing AM noise^36^, the posterior aspect of superior temporal gyrus for disrupted vowels_28_, and wider brain networks including the superior temporal lobe in the case of missed phonemes^24,37^. In addition, mixed evidence points to a basis for restoration in terms of endogenous modulations to boundary encoding: on the one hand, the search for differential onset responses to noise when under restoration, indexing alternative encoding, has yielded negative results so far^27,28^; on the other, induced narrow-band (3-4 Hz) desynchronizations that are restoration-specific, and occur after gap onset, have been suggested by results from EEG^28,38^.

In this study the differential temporal boundary encoding under restoration was not specifically addressed^38^, but instead the emphasis was on the neural representation of the missing rhythm itself, via measures of evoked rhythmic MEG responses. While restoration of continuous tones has been observed for segments as long as 1.4 s^39^ behaviorally, to our knowledge this is the first investigation where cortical aSSRs are directly implicated in perceptual restoration, sustained in real time representing a temporal code. That neural phase information was not reliable, despite an apparent continuity of the rhythm, is consistent with behavioral analyses suggesting that listeners may not track phase information under illusory FM continuity^8,9^. An cortical EEG study by Vinnik and colleagues^27^ showed no change to neural spectral power sustained along noise gaps embedded in a 40 Hz AM context stimulus during restoration; on the other hand, it has been shown that changes to neural spectral power in brainstem responses may occur during restored pitch of a missing 800 Hz carrier tone^29^. It is possible that while gamma-rate acoustic modulations can be represented cortically with a temporal code^13,40^, they are also at rates that involve pitch quality – a representation of which implies substantially distinct cortical coding modes^41^ assisting restoration.

In other sensory modalities, some restorative phenomena may fall in the category of perceptual experience that does not represent the absence of a physical stimulus, but rather, an alternative interpretation based on additional contextual information, e.g., the case of illusory induction of perceived kinesthetic trajectories^42,43^, and of spatial contours in certain visual displays^44–46^. Context-sensitivity in general is considered a requisite for cortical predictive coding^47^, which in the case of hearing may depend on known priors regarding the sound temporal dynamics. A compelling example arises from missing, but highly expected, click-like sounds that generate auditory onset-like responses locked to the nominal time of delivery of the missed sound^48^. Additionally, long duration, rhythmic metric structures may produce endogenous neural locking to a subharmonic frequency of the actual acoustic beat when it has the potential to be perceived as the underlying rhythm, whether listeners are instructed to do so^49^, or passively listen in the absence of instruction^50^. Correspondingly, the data here show that with perceptual restoration of masked rhythm, endogenous representational differences may emerge as early as 0.6 s post masking, at the target rhythm. There is also activity at the first harmonic, 10 Hz, but there one cannot entirely rule out yet alternative explanations involving enhanced alpha activity^27^, since with increased alertness at some trials over others, a systematic differential in spontaneous alpha activity might be responsible^51^. For filling-in and rhythm-missed trials related to inattention, reduced vigilance might be expected to effectively increase alpha activity. We did not, however find this; instead, filling-in trials displayed a narrow-band 10 Hz power increase strongly concurrent with the target duration, therefore consistent with being a harmonic of the endogenous 5 Hz rhythm. Alpha-band related effects due to non-uniform attentional states should be investigated in future studies using rhythms whose first harmonics are not in the alpha band. Our data does not reject the possibility of spontaneous and temporally patterned cortical activity profiles influencing sensory processing, as in ongoing slow-wave activity that may interact with evoked signals as a temporally coordinated modulation of excitability across distributed cortical fields^52,53^.

Focus on analysis of endogenous activity may address circumstances under which the brain repairs certain temporal features of highly stereotyped sound. This is part of the general problem of determining what relationship does a neurally-instantiated representation of a missed pattern has with a template representation mapping to actual acoustic experience. Solutions may offer key insight into biologically-inspired applications dealing with incomplete information. In particular, the modulation studied here corresponds to the temporal scale of syllabic production in human speech^54^ and the slow temporal envelope of natural stimuli^55^, thus raising the question of whether similar restorative phenomena exist during sequences of inner or imagined speech, as well as during auditory hallucinations.

## Methods

### Participants

35 subjects (12 women, 25.7 ± 4.4 years of age) with no history of neurological disorder or metal implants participated in the study, and received monetary compensation proportional to the study duration (~ 2 hours). Experiment protocol was approved by the UMCP Institutional Review Board; informed written consent was obtained from each participant.

### Stimuli

Four template sound stimuli were constructed with MATLAB^®^ (MathWorks, Natick, United States), each consisting of ~15 minutes of a 1024 Hz tone frequency-modulated (FM) at 5 Hz with modulation range (log-sinusoidal) 512–2048 Hz and a 20% duty cycle^16^. 420 rhythmic probes were created by adding 1.24 s of noise to the basic stimulus, at pseudo-random times. Noise was generated de novo per probe, and spectrally matched to the FM but with random phase. A fixed signal-to-noise ratio value was chosen from the -4 to 4 dB range, per participant. 420 non-rhythmic type trials were additionally created in the same manner, except that the underlying FM was replaced with constant carrier frequency. Inter-probe time intervals were 1.6 s plus a discrete Poisson-distributed random delay (λ = 1.2 s); the exact onset time was rounded to a multiple of the stimulus period (0.2 s), so that all probe onset times kept constant phase with the main rhythm. Sound stimuli were delivered through Presentation^®^ (NeuroBehavioral Systems, Berkeley, United States), equalized to be approximately flat from 40–3000 Hz, at a sound pressure level ~ 70 dB. Sounds were transmitted via E-A-RTONE® 3A tubes (impedance 50 Ω) and E-A-RLINK® disposable foam intra-auricular ends (Etymotic Research, Elk Grove Village, United States) inserted in the ear canals.

### Experimental design

After a brief practice session, subjects were instructed to push one of a pair of buttons based on whether they detected a 5 Hz rhythm. In order of importance, participants were instructed to: (i) wait until probe ended before pressing the button, weighting accuracy over reaction time; (ii) respond only to the probe immediately presented;(iii) modify their choice by pressing the other button only if certain and still before the next trial. Trials that did not meet the requirements, and corrected trials, were excluded (median 6.8% and 1.3% of trials respectively). To avoid transient cortical dynamics associated with motor response execution^56^, trials beginning less than 250 ms from the previous response were also excluded (median 6.3% of trials). To more evenly distribute the proportion of correct answers across participants, the masker signal-to-noise ratio (SNR) was fixed in advance, from on of 0, ±1, ±2 or ±4 dB. Silent films were presented concurrently, which subjects were instructed to watch.

### Data recording

MEG data were collected with a 160-channel system (Kanazawa Technology Institute, Kanazawa, Japan) inside a magnetically-shielded room (Yokogawa Electric Corporation, Musashino, Japan). Sensors (15.5 mm diameter) were uniformly distributed inside a liquid-He Dewar, spaced ~25 mm apart. Sensors were configured as first-order axial gradiometers with 50 mm separation and sensitivity > 5 fT·Hz-1/2 in the white noise region (> 1 KHz). Three of the 160 sensors were magnetometers employed as environment reference channels. A 1 Hz high-pass filter, 200 Hz low-pass filter, and 60 Hz notch filter were applied before sampling at 1 KHz. Participants lay supine inside the magnetically shielded room under soft lighting, and were asked to minimize movement, particularly of the head. Every session had four experimental blocks. In the case of seven participants, the experiment had to be suspended early due to time constraints (mean 89% completion in these participants, minimum 75%); for one participant only 2 blocks out of 4 were recorded due to transfer failure. Two participants requested pauses during a block, which was terminated and later repeated in whole.

### Data processing

A 1-30 Hz band-pass third order elliptic filter with at most 1 dB ripple and 20 dB stopband attenuation was applied and noise sources were removed as follows. *Environment noise*. Time-shifted principal component analysis^57^ (TS-PCA) was applied to remove environmental noise, using the three reference magnetometers (*Nlags* = 43). *Sensor-specific noise*. Sensor-generated sources unrelated to brain activity were subtracted using sensor noise suppression (SNS)^58^. *Spatial filtering*. A per-participant data-driven model was used to synthesize spatial filters from the responses to the unmasked rhythmic sound stimulus via denoising spatial separation (DSS)^59^. The responses were structured as a matrix of dimensions *T* x *N* x *K*; where *T* is the number of samples (=1400), *N* is the number of usable recording segments (average=514.3), and *K* the number of active sensors (average=156.8).This spatial filter selects for the most reproducible aSSR component over trials, generating a single virtual sensor used in the remaining analysis.

### Data analysis

Trials were classified a posteriori, according to subjects’ reports, into one of four groups: rhythmic-trial perceived such (‘driven’) or as not as non-rhythmic (‘missed’);non-rhythmic trial perceived as such (‘absent’), or as rhythmic (‘filling-in’). Time-frequency analysis used a Morlet wavelet transform with 0.2 s scale, permitting estimation of spectral evoked power at the bandwidth of experimental interest (5 Hz). For evoked power and ITPC contrasts, statistical clusters were found during which there were significant differences across experiment conditions according to non-parametric permutation tests^60^. A measure of neural discriminability’, 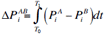is defined as the area between two evoked power curves *P* obtained each at conditions *A* and *B* for the *i*-th subject, and computed over a fixed time interval (*T*_*0*_ = 0.58 s and *T*_*1*_ =1.2 s post noise onset on average), as defined by statistical clusters of significance found at the group level for the given contrast *AB*. Measures for shifts in ITPC were computed in similar way. Perceptual sensitivity of a subject in detection is given by *d*-prime analysis^19^, 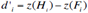 where for each subject *i, H* the fraction of rhythmic probes labeled rhythmic, and *Fi* the fraction of non-rhythmic probes labeled rhythmic, undergo a z-transformation.^61^

To investigate whether the observed pattern of percept-specific differences was due to unintended acoustical or statistical properties in the stimulus constructs, stimulus probes were analyzed a posteriori. No significant differences were found in stimulus temporal modulations when partitioned by percept, within rhythmic (*p*=0.85) nor non-rhythmic (*p*=0.84) probes (paired-sample *t*-tests, Supplementary Fig. S3).

Subjects’ reported percepts corresponded to the physical acoustics (presence or absence of rhythm) approximately 5 times as often as not, resulting in data pools with differing signal-to-noise ratio improvement from averaging. Therefore inter-trial phase coherence measures included bias correction^62^ as small sample sizes are especially prone to bias. The unbiased estimator is based on the squared ITPC (also defined as squared ‘modified resultant length’^62^), which may be negative after estimated bias subtraction. To investigate the possibility of related biases in the rhythmic evoked power measures, post hoc two-sided non-parametric permutation tests were performed by collecting, for each subject, all trials from the two conditions to be compared, and instantiating resampled of partitions of fixed size (original sample sizes per subject); the group-level test statistic obtained in the actual partition was then contrasted against those obtained at group level across the distribution of resampled instances. Using the 5 Hz evoked power difference between conditions in the same intervals of significance, it was found that responses to non-rhythmic probes show significantly greater power when reported perceived as rhythmic versus non-rhythmic (0.56 to 1.19 s; *p*=0.007); a similar result held for responses to rhythmic probes, which also show significantly greater power when reported perceived as rhythmic versus non-rhythmic (1.04 to 1.15 s; *p*=0.034). Potential systematic differences resulting from the per-subject signal-to-noise (SNR) ratio were also investigated, but no evidence was found of differences, neurally (*ρ*=0.10, *p*=0.57) or behaviorally (*ρ=*0.33, *p*=0.054). One participant was excluded from the analysis due to zero reported perceptual differences from the acoustics.

### Data availability

All relevant data are available to all interested parties in a public repository.

## Acknowledgements

This study was funded by the National Institutes of Health (R01-DC-00843, R01-DC-014085). We thank support to FCC by the Mexican National Council of Science and Technology through its graduate scholarship program. We thank Natalia Lapinskaya for excellent technical assistance, and Richard Payne, Matthew Goupell, Jonathan Fritz, Ellen Lau, Daniel Butts and Behtash Babadi for discussions.

## Author Contributions

FCC conceived, designed, and performed the experiments, analyzed the data, and prepared the manuscript figures. JZS supervised the research. Both authors wrote the manuscript text.

